# I will teach you here or there, I will try to teach you anywhere: perceived supports and barriers for emergency remote teaching during COVID-19 pandemic

**DOI:** 10.1101/2021.06.21.449058

**Authors:** Cristine Donham, Hillary A. Barron, Jourjina Alkhouri, Maya Changaran Kumarath, Wesley Alejandro, Erik Menke, Petra Kranzfelder

## Abstract

**Background:** Due to the COVID-19 pandemic, many universities moved to emergency remote teaching (ERT). This allowed institutions to continue their instruction despite not being in person. However, ERT is not without consequences. For example, students may have inadequate technological supports, such as reliable internet and computers. Students may also have poor learning environments at home and may need to find added employment to support their families. Additionally, there were consequences to faculty. It has been shown that female instructors are more disproportionately impacted in terms of mental health issues and increased domestic labor.

This research aims to investigate instructors’ and students’ perceptions of their transition to ERT. Specifically, we wanted to:

1. Identify supports and barriers during the transition to ERT
2. Compare instructors experiences with the student experiences
3. Explore these supports and barriers within the context of *social presence, teaching presence*, and/or *cognitive presence* during ERT as well as how these supports and barriers relate to *scaffolding* in emergency remote courses

**Design:** Using grounded theory techniques, we applied two-cycle, qualitative analyses to assess the instructor transcripts. In first-cycle analysis, we used open coding to develop initial ideas from the data. We then used second cycle coding to generate categories with definitions and criteria agreed upon during discussion-based consensus building. Finally, these categories and descriptions were used to code student survey data.

**Analyses/Interpretations:** Instructors identified twice as many barriers as supports in their teaching during the transition to ERT and identified casual and formal conversations with colleagues as valuable supports. Emerging categories for barriers consisted of academic integrity concerns as well as technological difficulties. Similarly, students identified more barriers than supports in their learning during the transition to ERT. More specifically, students described pre-existing course structure, classroom technology, and community as best supporting their learning. Barriers that challenged student learning included classroom environment, student availability, and student emotion and comfort.

**Contribution:** Together, this research will help us understand supports and barriers to teaching and learning during the transition to ERT. This understanding can help us better plan and prepare for future emergencies, particularly at MSIs, where improved communication and increased access to resources for both students and instructors are key.

## INTRODUCTION

In the middle of the Spring 2020 academic term, many institutions of higher education were forced to move all instruction online. The term “pandemic pedagogy” was quickly coined as educators, many of whom had never taught online or remotely, scrambled to come up with effective ways to teach their courses (Schwartzman, 2020). Although moving to remote instruction can enable flexibility of teaching and learning (Daymont et al., 2011), the speed at which instructors and students were expected to move to remote instruction was unprecedented. Therefore, it is important to distinguish this quick transition to remote teaching, or *emergency remote teaching* (ERT), from the traditional *online teaching*. Here, we use the term *online teaching* to refer to traditional online teaching (i.e., teaching online during non-pandemic times), and have adopted Charles Hodges’s definition of ERT as a temporary shift of instructional delivery to an alternate delivery mode due to crisis circumstances (Hodges et al., 2020). ERT is characterized by improvised, quick solutions in less-than-ideal circumstances, and it was the best solution most universities had to academic learning. This is different from traditional *online teaching* where instructors are intentionally designing a course to be implemented and delivered online, a delivery mode that has been studied for decades (Bender, 2012; Lewis & Abdul-Hamid, 2006; Oliver, 1999; Young, 2006). There are numerous research studies, theories, models, and evaluation criteria created for traditional *online teaching* (Oliver, 2000; Ouyang & Scharber, 2018; Shelton & Hayne, 2017). Studies have shown that effective online learning stems from careful instructional design, planning, and using a systematic model for the development (Branch & Kopcha, 2014). This careful design process was likely to be absent in most ERT shifts due to lack of time and experience necessary for instructors to carefully design their course for online purposes.

In addition to a shortage of time and experience, the move to ERT introduced a variety of issues that instructors and students had not faced during in-person teaching, such as lack of communication (Gelles et al., 2020). Previous research on emergency teaching during Hurricane Katrina in 2005 showed that virtual student-to-student interactions and remote class dialogues created opportunities for students to provide mental and emotional support for each other (Lorenzo, 2008). Other issues with ERT include navigating the course in a new manner, finding new ways to implement formative assessment, communicating with students in a fair and equitable manner, monitoring academic integrity, and managing everything through a remote platform (Brooks & Grajek, 2020; Johnson et al., 2020). Additionally, many students moved home and were expected to attend college from home while being quarantined with their family. Students needed to learn how to navigate a new learning platform and adjust to new patterns, all while experiencing loss of social interactions. Shay and Pohan 2021 described how “many low-income, first-generation students also struggle with a lack of quiet workspaces, the absence of internet and other technological tools, housing and/or food insecurity, and the added responsibilities associated with being at home and helping the family (e.g., employment, care, etc.)” (Shay & Pohan, 2021). Stress, anxiety, and traumatic events contributed to students’ cognitive load, - the demand or burden on one’s working memory - making the focus on learning more challenging for students (Shay & Pohan, 2021). Finally, high-quality internet is an enabling technology for ERT, and in the US approximately 21 million people, or 6.5% of the population, do not have broadband internet access (Chavez, 2020; Commission, 2018). This lack of access to technology during ERT created a digital divide in higher education for both instructors and students. Additionally, despite most institutions transitioning to ERT, there is a lack of literature examining ERT experiences in higher education, particularly for college science, technology, engineering, and mathematics (STEM) education (Johnson et al., 2020). Since COVID forced ERT began, there have been some new literature examining a wide variety of impacts on students and instructors (Affouneh et al., 2020; Bozkurt & Sharma, 2020; Iglesias-Pradas et al., 2021; Karakaya, 2021; Whalen, 2020). Despite this new literature, there is a need to fill this gap in literature by examining what helped and hindered both instructors and students during their transition to ERT. We aim to help fill this gap by studying supports and barriers due to COVID-19 ERT at a minority serving institution.

The University of California (UC), Merced is in a unique situation to study the rapid transition to ERT. While it is a member of a well-resourced university system, its student population is highly diverse and similar to student populations that are typically understudied in education (Council, 2012; Kanim & Cid, 2020). For example, during the 2019-2020 academic year the UC Merced undergraduate population was 74% first-generation, 54% Hispanic (and only 9% non-Hispanic white), and 64% Pell-Grant eligible. This student population is representative of populations across the country that are disproportionately affected by COVID-19, as they are vulnerable to living in persistent poverty, holding unstable or uncertain employment, and without stable housing (Burke, 2020).

Additionally, UC Merced’s faculty population is younger (83% of UC Merced STEM faculty under age 55, compared to 62% across the UC system) and more likely to be a woman (35% of UC Merced STEM faculty are women, compared to 24% across the UC system) than faculty in the UC system as a whole (University of California, 2020a) (University of California, 2020b). Like the student population, the faculty population is more likely to suffer personally and professionally, from the negative impacts of the COVID-19 pandemic than the broader UC population. Female instructors in general, are thought to be more disproportionately impacted in terms of mental health issues and increased domestic labor due to the pandemic (Brooks & Grajek, 2020; Donner, 2020; Krentz et al., 2020). Because of these factors, we wanted to understand the supports and barriers experienced by UC Merced instructors and students during the transition to ERT.

### Theoretical Frameworks

The work we present here is guided by the *Community of Inquiry* (COI) framework (D Randy Garrison et al., 2010). The COI framework is a collaborative-constructivist process model that describes the essential elements of successful online higher-education learning experiences (Garrison, 2016; Garrison et al., 2000). The COI framework has been widely used when investigating traditional *online learning* as it is grounded in a social constructivist approach to learning (Garrison et al., 1999) (e.g., Arbaugh et al., 2008; Castellanos-Reyes, 2020) (Piaget, 1976). Additionally drawing from this social constructivist lens, learners co-construct knowledge by engaging in actions that elicit and validate their sociocultural identities (Dewey, 1986; Vygotsky, 1978). Vygotsky contended that socially-situated learning, accompanied by scaffolding, led to stronger outcomes in both knowledge development and retrieval (Dewey, 1986; Vygotsky, 1978). In addition to COI, we utilized the scaffolding framework as a collaborative-constructivist process model from which we examined how instructors have adapted to remote pedagogy and the supports and barriers that instructors and students found helped them or hindered them. Scaffolding refers to the help or guidance from a more competent peer or mentor that allows students to work within, and then move beyond, the zone of proximal development (ZPD) (Wass et al., 2011).

#### Community of Inquiry

Generally, COI is a process model of online learning which views the online educational experiences as arising from the interaction of three elements: *teaching presence, social presence*, and *cognitive presence*. In the COI framework, *teaching presence*, is defined as the design, facilitation, and direction of cognitive and social processes to realize teacher-defined learning outcomes. This element has three components: 1) instructional design and organization (e.g., instructor provides clear instructions on how to achieve course learning outcomes); 2) facilitating discourse (e.g., instructor helps to keep course participants engaged and participating in productive dialogue); and 3) direct instruction (e.g., instructor presents useful examples that allows students to better understand course content) (Anderson et al., 2001b) (Figure 1). Research has shown that *teaching presence* is important for successful online learning and strongly correlates with student satisfaction, perceived learning, and sense of community (Kucuk & Richardson, 2019; Liu, 2019; Manwaring et al., 2017; Soffer & Cohen, 2019). Shea and Bidjerano (2008) found that the quality of *teaching presence* and *social presence* reported by learners in online courses could predict learning represented by the *cognitive presence*. Additionally, D.R. Garrison et al. (2010) found that student perceptions of *teaching presence* predicted significant direct effect on perceptions of *cognitive presence*, while *social presence* had an indirect or mediating effect on *cognitive presence*. Therefore, *teaching presence* has been shown to be central in creating quality online learning experiences.

**Figure 1.**
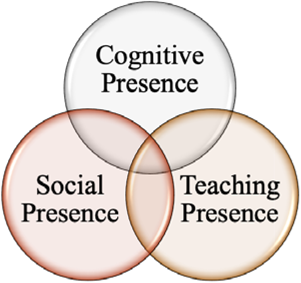
Community of Inquiry (COI). The COI theoretical framework represents a process of creating a deep and meaningful, collaborative-constructivist, learning experience through the development of three interdependent elements – social, cognitive, and teaching presence. Taken from Anderson, Rourke, Garrison, & Archer, 2001.

The second element in the COI framework, *social presence*, is defined as the ability for learners to be perceived as “real people.” It is primarily focused on affective expression (e.g., students perceived online or web-based communication as an excellent medium for social interaction), open communication (e.g., students felt comfortable interacting with other course participants), and group cohesion (students perceived online discussions as helping them develop a sense of collaboration). Research on *social presence* has demonstrated a strong relationship between *social presence* and student learning outcomes (d’Alessio et al., 2019; Hwang & Arbaugh, 2006; Richardson et al., 2017). Additionally, research has shown that activities that cultivate *social presence* also enhance learner satisfaction (Richardson et al., 2017), and that interaction and engagement through active learning is necessary for students to feel as they are dealing with real people, that they belong in some way to a group of learners, and that they are involved in sharing, negotiating, arguing, and discussing (Wang, 2008). In general, online learning environments should be active, allow student to construct their own knowledge, make effective use of collaborative and cooperative learning methods, and be meaningful to students while promoting *social presence* and community (Ally, 2004).

Finally, the third element, *cognitive presence*, is defined as the extent to which learners can construct and confirm meaning through sustained reflection and discourse. It is rooted in social constructivism, which is a robust theory proposing that people’s learning is shaped by cultural context, conversation, and collaboration (Dewey, 1986; Vygotsky, 1978). *Cognitive presence* has four components: 1) triggering event (e.g., students felt motivated to explore content related questions); 2) exploration (e.g., students used a variety of information sources to explore problems posed in the course); 3) integration (e.g., learning activities helped students construction explanations/solutions); and 4) resolution (e.g., students applied the knowledge created in the course to their work and non-class related activities). Of the three elements in COI, *cognitive presence* has been identified as the most difficult to study as well as the most challenging to develop and sustain in online courses (Garrison & Cleveland-Innes, 2005). This difficulty arises from the fact that *cognitive presence* contains inputs (the triggering event), processes (exploration and integration), and outputs (resolution) that can be hard to measure or observe, whereas the other two elements consist of processes that can be more easily observed (Garrison & Arbaugh, 2007).

While the COI framework has been tested for online learning and K-12 teachers during ERT (Whittle et al., 2020), it has only recently been used for ERT in undergraduate STEM classrooms at a Minority-Serving Institute (MSI). For example, Reinholz *et al*. used the COI framework to study how the nature of student participation changed in moving from face-to-face to synchronous online learning environments at an Hispanic serving institute (HSI) (Reinholz et al., 2020) and Eriksen *et al*. studied how student perceptions of *social presence, cognitive presence*, and *teaching presence* online due to COVID-19 were influenced by student demographics (Erickson & Wattiaux, 2021).

#### Scaffolding in distance learning

In 1976, Wood, Bruner and Ross introduced the term *scaffolding* (Wood et al., 1976) and many researchers and educators have used the concept of scaffolding to describe instructor roles as more knowledgeable peers for guiding student learning and development (Hammond, 2001; Stone, 1998; Wells, 1999). Scaffolding has been interpreted in a wide sense as “a form of support for development and learning” (Rasmussen, 2001, p570). Alternatively, it can be used as an umbrella metaphor to describe the way that teachers supply students with the tools necessary to learn (Jacobs, 2001).

While there are different approaches in the literature on how *scaffolding* may or may not be intertwined with Vygotsky’s ZPD (Vygotsky, 1980), we used Wells’s argument that *scaffolding* can be a direct application and operationalization of ZPD (Berk, 2003; Duchesne & McMaugh, 2018; Wells, 1999). Wells identified three features that characterize educational *scaffolding:* 1.) The essentially dialogic nature of the discourse in which knowledge is co-constructed (dialog); 2.) the significance of the kind of activity in which knowing is embedded (activity); and 3.) the role of artifacts that mediate knowing (artifacts) (Wells, 1999, p.127). Furthermore, the relationship between classroom challenge and support is important in *scaffolding* (Hammond & Gibbons, 2005). Hammond and Gibbons (2005) found that highly supportive, but minimally challenging environments may be too easy to elicit growth in knowledge, whereas experiences that are highly challenging but lack sufficient support will likely result in failure. This becomes an important aspect of *scaffolding* to consider as we examine how instructors and students experienced the switch to ERT.

Therefore, we had three main objectives:

1. Identify supports and barriers experienced by instructors and students during the transition to ERT.
2. Compare the instructor experience with the student experience.
3. Explore these supports and barriers within the context of *social presence, teaching presence*, and/or *cognitive presence* in the remote teaching environment as well as how these supports and barriers relate to *scaffolding* in emergency remote courses (Figure 2).

**Figure 2.**
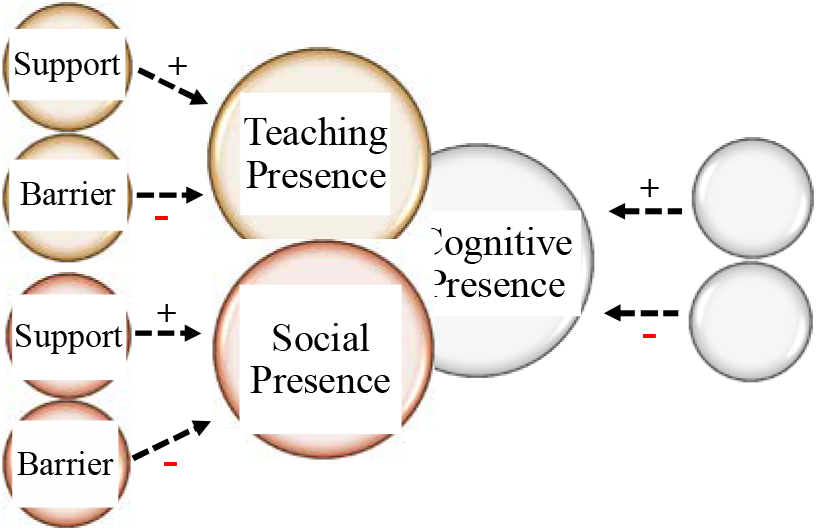
The Community of Inquiry. Each support and barrier influenced its designated presence/attribute and each presence/attribute ultimately led to a specific *cognitive presence* for instructors and students. (+) indicate positive influences in each presence or attribute and (-) indicates negative influences.

## METHODS

### Recruitment

The study was approved by the Human Subjects Committee of the University of California Merced’s Institutional Review Board (IRB) (Protocol ID UCM2020-3).

#### Instructor

In May 2020, we sent an initial recruitment email, via departmental list serves and individual email addresses, to tenure-track faculty, adjunct faculty, and teaching assistants who were teaching STEM courses at UC Merced. The initial email included the purpose of the study, procedures, benefits, IRB approval, potential dissemination of the results, question information, and contact information. Instructor recruitment occurred approximately two months after the institutional switch to ERT.

We chose the instructors in this study due to their ongoing involvement in a larger institutional study on assessing teaching and discourse practices in college STEM classrooms. This larger study focused on instructors who 1) taught either a lower or upper division undergraduate or graduate STEM course and 2) taught a lecture or laboratory course. To recruit participants for this study on the transition to ERT, we applied the same requirements (1 and 2) plus two additional criteria: 3) taught a remote course via synchronous instruction (excluded in-person and asynchronous instruction), and 4) taught in Spring 2020.

#### Student

During the instructor recruitment process, all instructors were asked if they would be willing to invite their students to a 30-minute in-class or out-of-class group interview. Out of 31 instructors that agreed to participate in the instructor interviews, 22 agreed to inviting their students to these interviews (71% participation rate). Students from five courses attended and agreed to participate in these interviews (16% participation rate).

### Population

We interviewed 31 instructors and 69 students at a mid-sized, public, research-intensive university designated as an MSI. Instructors taught biology, chemistry, engineering, mathematics, and physics courses, while students were in biology, chemistry, and physics courses (Table 1). Gender and ethnically appropriate pseudonyms were created for instructors and gender-neutral pseudonyms were created for students to de-identify participants and retain their privacy and confidentiality.

**Table 1.** Demographics of instructors (Faculty and TAs) (*n*=31) and students (n=69)

| **STEM discipline** | *Faculty* | *TA* | *Student* |
| --- | --- | --- | --- |
| Biology | 10 | 5 | 1 |
| Chemistry | 4 | 4 | 31 |
| Engineering | 2 | 2 | 0 |
| Mathematics | 2 | 1 | 0 |
| Physics | 1 | 0 | 37 |

### Data collection

#### Instructor

Instructor interviews were carried out via Zoom between May 11 and May 27, 2020. The semi-structured interviews ranged in length from 10-90 minutes, where the participants were asked six questions regarding perceived supports and barriers, changes in interactions with students and instructors, pedagogical changes made or planned, and potential future supports. This manuscript will focus only on perceived supports and barriers (Table 2). Instructors were not interviewed by anyone from their own department (i.e., biology faculty interviewed chemistry faculty). Four of the authors conducted the 31 interviews. After the interviews were completed, they were transcribed using a clean verbatim transcription service.

**Table 2.**
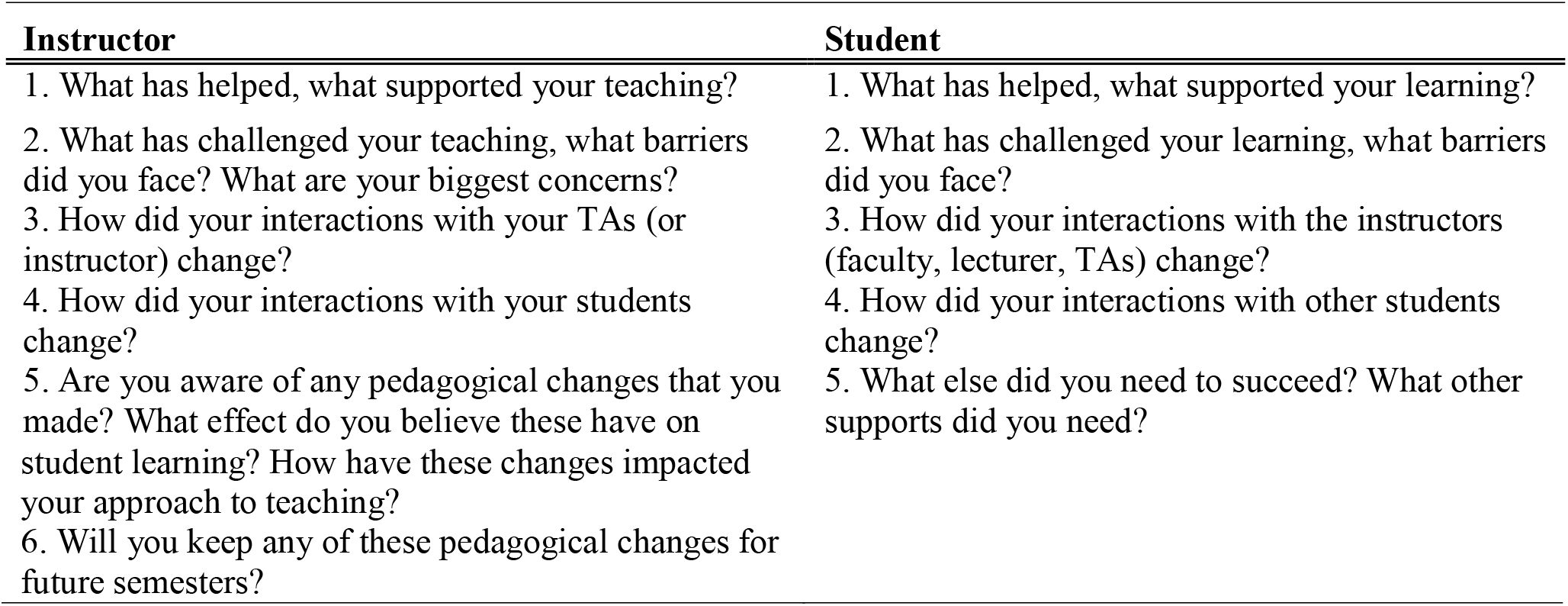
Interview questions

#### Student

Student surveys and group interviews were carried out via Zoom during the final week of instruction (between May 5 and May 8, 2020) by three Students Assessing Teaching and Learning (SATAL) interns, an assessment team of trained undergraduate students out of UC Merced’s Center for Teaching and Learning (CETL) (Signorini & Pohan, 2019). Individual surveys and group interviews lasted about 30 minutes during discussion sections, where the participants were asked five questions regarding perceived supports and barriers, changes in interactions with instructors and other students, and potential future supports. This manuscript will focus only on perceived supports and barriers from the student surveys (Table 2).

### Data analyses

#### Instructor interview coding

Grounded theory techniques (Charmaz, 2006; Strauss & Corbin, 1998) in qualitative analysis employ a rigorous, iterative process of examining the properties and dimensions of data in order to create a holistic understanding of a process or phenomenon (Charmaz, 2006; Corbin & Strauss, 1990; Miles et al., 2018). Drawing from grounded theory methods, we used two cycle qualitative analysis (Miles et al., 2018) to explore instructor interview transcripts in a section-by-section fashion (Figure 3).

**Figure 3.**
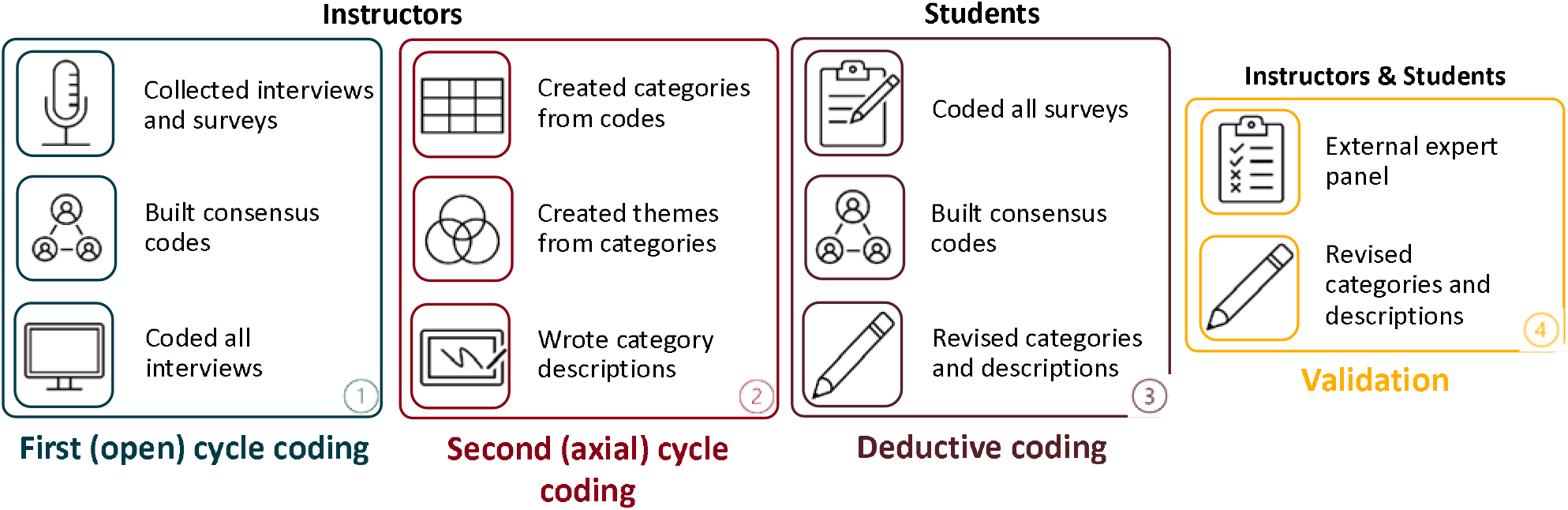
Data analysis. Overall scheme depicting methods used. First cycle coding was completed for instructor interviews followed by second cycle coding where categories and themes were generated from initial codes. Following instructor data analysis, student survey data was coded by authors using categories generated previously. Following instructor and student analysis, an external expert panel was used to validate categories and descriptions. The color blue represents first or open cycle coding, the color red represents second cycle coding, the dark red represents deductive coding with student data, and the color yellow represents study validation.

##### First (open) cycle coding

First-cycle qualitative coding allows researchers to gain a comprehensive and integrated view of a dataset (Miles et al., 2018). It is intentionally cyclical, such that a code generated during the first cycle is not meant to be a static assessment. Rather, fluidity is essential. As we engaged in first cycle analysis, we used open coding to look holistically across all the data, and to identify repeating indicators of instructors’ perceptions of teaching during the COVID-19 pandemic. Strauss and Corbin (1998) describe open coding as a process of examining properties and dimensions that exist within the data, allowing the researcher to identify unique and discrete aspects (Miles et al., 2018). To do this, we looked across all instructor transcripts and began assigning codes that indicated how instructors perceived supports and barriers while they conducted ERT. Table 3 demonstrates what first cycle, open coding looked like for one of the two researchers (CD and EM).

**Table 3.** First (open) cycle coding

| ID | Dialogue | CD initial code |
| --- | --- | --- |
| Interviewer | *What would you say was the biggest concern for the transition? I know you said the communication with students was bad.* |  |
| Samuel | *For what I saw on the students' side, access to technology was sometimes difficult. I didn't have too many examples, but a recent thing is one student, their parents have both lost their job, and so, the student has to start a job on their own, working as a fruit picker in the farm, and not being able to have the availability for exam minutes, before. So, asking for an exception to take the exam.* | **Barrier:** lack of student access to technology; **Barrier:** Student obligations and pressures |

Prior to moving into the second cycle of analysis, we engaged in consensus-building with five researchers (CD, EM, PK, WA, and HB). Consensus building in qualitative analysis is a critical measure of ensuring validity and trustworthiness (Corbin & Strauss, 2014). To address inter-coder consistency, we independently coded approximately 10% of all transcripts, and then we used discussion-based consensus building to address discrepancies in codes. Saldaña (2015) describes this process as *interpretive convergence*, specifically useful in qualitative analysis where dynamic interpretations of data are paramount, as opposed to seeking statistical significance in quantitative methodologies. In our efforts to converge toward common codes, we discussed both our individual open codes as well as our analytic memos. Analytic memos in qualitative analysis serve as a researcher’s dialogue, both with themselves and each other, about what the codes mean to them as they are coding (Charmaz, 2006; Corbin & Strauss, 2014; Miles et al., 2018; Stake, 2005). Table 4 depicts open codes and analytic memos during one of our consensus coding sessions. It shows five of the author’s initial codes, generated individually, and the consensus code generated as a group after discussion.

**Table 4.** Consensus coding

| **Dialogue** |  |  |  |  |  |
| --- | --- | --- | --- | --- | --- |
| *So one of the challenges there is to prevent students from communicating, but we couldn't do it with a large class just like we had with Bio 2. One section was over 200 students. So what I had to do was use a trick. 'Cause we were using breakout rooms. Maybe, I'll come back to that in a different question. But to finish off your question here, I've tried at Stanislaus to have 150 students on camera with their microphones, and it worked. Although, three students did drop and have to reconnect. So I found that useful to have pictures of that—all the students taking the exam using either their screen camera, or if they didn't have a working computer camera, I made them put their phone next to them and themselves so I could see what their eyes were looking at. --- Kailish* |  |  |  |  |  |
| **Initial codes** |  |  |  |  |  |
| **CD** | **EM** | **PK** | **WA** | **HB** | **Consensus** |
| Challenge to academic integrity; made students use camera | Student academic integrity; preventing cheating | Student videos are possible and preferred; instructor monitors students during in-lecture, timed exams to reduce academic misconduct | Previous experience; prefers students to have webcams on | Couldn't prevent students from communicating | **Barrier:** Monitoring students during synchronous exams to prevent cheating |
| **Analytic Memos** |  |  |  |  |  |
| **CD** | **EM** | **PK** | **WA** | **HB** |  |
|  |  | Student academic conduct during exams seems like a key challenge for instructors teaching with ERT | Instructor has experience with having a large number of participants in a video call on webcams | Others have mentioned academic honesty/integrity in this section too |  |

##### Second (axial) cycle analysis

The aim of second cycle analysis is to find linkages between the discrete parts that were earlier identified in open coding (Miles et al., 2018), in order to find “broader categories, themes, theories, and/or assertions” (p. 234). Essentially, this involves looking for similarities and differences across the previously identified properties and dimensions of the dataset (Strauss & Corbin, 1998). We engaged in this process of creating relational categories through axial coding. Table 5 and Table 6 depicts our process of axial coding.

**Table 5.**
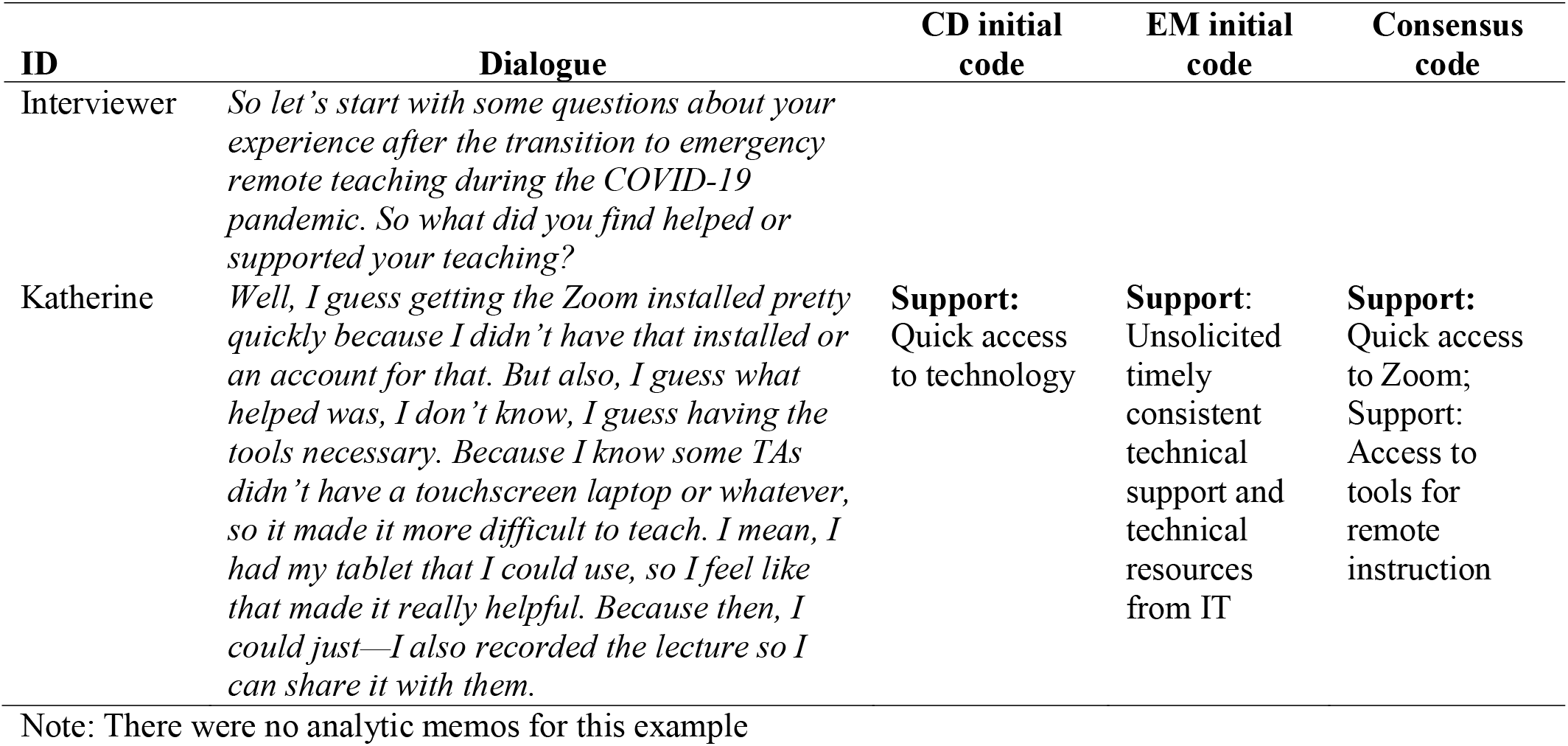
Second (axial) cycle consensus coding with support example

| ID | Dialogue | CD initial code | EM initial code | Consensus code |
| --- | --- | --- | --- | --- |
| Interviewer | *So let's start with some questions about your experience after the transition to emergency remote teaching during the COVID-19 pandemic. So what did you find helped or supported your teaching?* |  |  |  |
| Katherine | *Well, I guess getting the Zoom installed pretty quickly because I didn't have that installed or an account for that. But also, I guess what helped was, I don't know, I guess having the tools necessary. Because I know some TAs didn't have a touchscreen laptop or whatever, so it made it more difficult to teach. I mean, I had my tablet that I could use, so I feel like that made it really helpful. Because then, I could just—I also recorded the lecture so I can share it with them.* | **Support:**<br>Quick access to technology | **Support:**<br>Unsolicited timely consistent technical support and technical resources from IT | **Support:**<br>Quick access to Zoom; Support: Access to tools for remote instruction |

Note: There were no analytic memos for this example

**Table 6.** Second (axial) cycle consensus coding with barrier example

| ID | Dialogue | CD initial code | EM initial code | Consensus code |
| --- | --- | --- | --- | --- |
| Interviewer | *What would you say was the biggest concern for the transition? I know you said the communication with students was bad.* |  |  |  |
| Samuel | *For what I saw on the students' side, access to technology was sometimes difficult. I didn't have too many examples, but a recent thing is one student, their parents have both lost their job, and so, the student has to start a job on their own, working as a fruit picker in the farm, and not being able to have the availability for exam minutes, before. So, asking for an exception to take the exam.* | **Barrier:** Lack of student access to technology;<br>**Barrier:** Student obligations and pressures | **Barrier:** Student access to technology;<br>**Barrier:** Student obligations and pressures | **Barrier:** Lack of student access to technology;<br>**Barrier:** Student obligations and pressures |

Note: There were no analytic memos for this example

Four researchers (CD, EM, PK, and HB) worked together to make descriptions for supports and barriers categories. The following categories and descriptions emerged from the data (Supplemental Table 1). Support categories included: prior experience, timing, technology for remote teaching, community, help with technology, socio-emotional factors, teacher beliefs, working from home, help with teaching, course attributes, student comfort interacting online, and reducing cognitive load. Barrier categories included: communication difficulties, time management, instructor teaching inexperience, instructor technology issues, teaching and learning resources, student integrity, administrative issues, student presence and participation, student emotion and comfort, student technical issues, assessment difficulties, instructor emotion, responsibility and workload, and instructional space.

#### Student deductive coding

A deductive coding approach was used to identify the various student support and barrier categories generated from coding the instructor interviews (Table 2). First, two researchers (PK and MCK) independently coded the student responses using the 12 support categories (Table 9) or 14 barrier categories (Table 10) to get initial codes (Table 7). Each student response was one to three sentences in length and could contain more than one category.

**Table 7.**
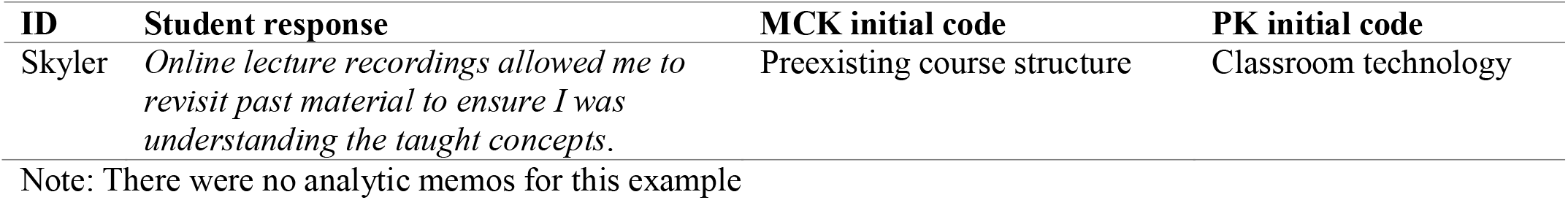
Individual coding with support example

Next, PK and MCK met to discuss their categories until reaching 100% consensus (Table 8). When coding student responses, authors noticed that the working descriptions of some categories would only fit the instructor perspective and needed to be redefined to be used for student coding. Authors then met to discuss which descriptions should be changed and rewrote them so that the codes could be used for both students and instructors alike.

**Table 8.**
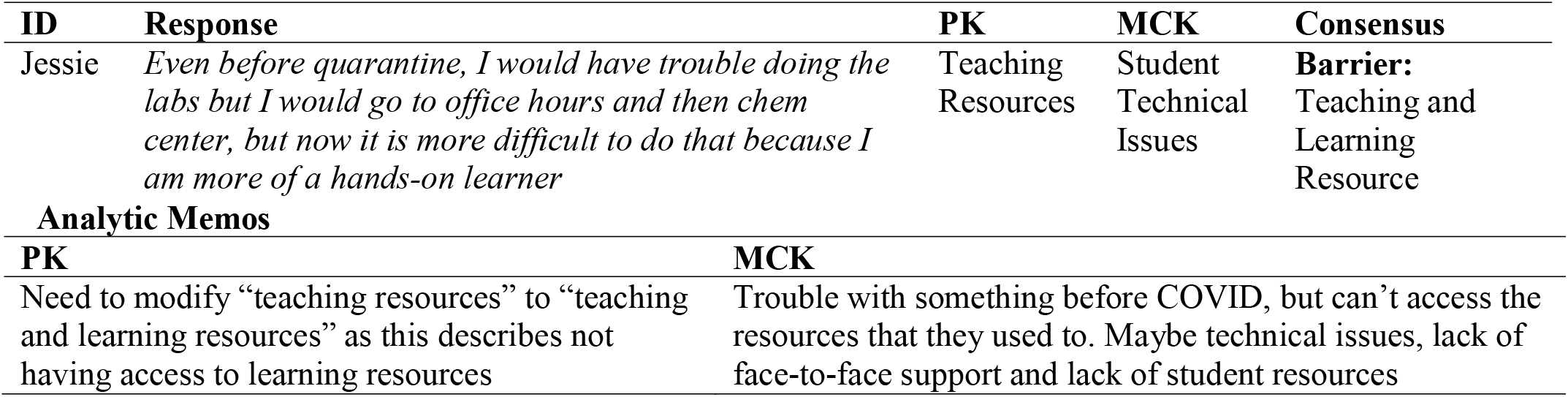
Consensus coding with barrier example

#### Validation

Following student coding using the categories and agreed upon descriptions, we brought the categories, themes, and descriptions of categories to an expert feedback panel of five. The expert feedback panel was made-up of STEM educators (both biology and chemistry), discipline-based education researchers (DBER), and learning scientists at a research-intensive institution unrelated to the one in this study. This expertise allowed the panel to provide valuable feedback on category descriptions.

The formatting for the feedback was organized in two parts. For the first part, the first author (CD) presented the support instructor and student themes, categories and their descriptions along with examples (Supplemental Table 1). Expert feedback panelists were then allowed to ask questions prior to a short content validation. This content validation consisted of splitting the panelists into two groups and providing each group with the same three *support* quotes representative of categories in each of the three themes (Supplemental Table 3 and 4). The panelists were given all the support categories and descriptions and were asked to match the quote with a category and provide justification via notes. Additionally, authors CD, PK and JA were present in both groups to take notes on feedback. After each group had finished choosing categories for the quotes, everyone met back in one group and the answers were discussed. Comments and feedback are reported in Supplemental Table 3 and 4. Following this discussion, the *barriers*, their descriptions, and themes were then presented by CD to the expert feedback panel (Supplemental Table 2). The same two groups were then created, and panelists were again given the list of barriers and their descriptions and asked to decide which category fit each quote best. After each group had finished all comments and questions were recorded (Supplemental Table 5 and 6). Again, CD, PK, and JA were present in both groups to take notes on feedback.

Following the expert feedback panel, all coders met and discussed the feedback. This discussion allowed changes from version 2 to version 3 for categories and descriptions (Supplemental Table 7 and 8). These are the final categories and descriptions presented in this manuscript.

## RESULTS

The findings presented here are separated into the two primary areas of study, supports and barriers. We identified 134 unique support codes which led to the development of 12 categories, and, ultimately, three support themes: 1) tools and support for class content, 2) mental & emotional support, and 3) pre-existing supports. A total of 203 unique barrier codes were identified, which led to the development of 14 categories from which the following three barrier themes emerged: 1) structures getting in the way, 2) spending more time and effort, and 3) affective issues.

### Instructor supports

Instructors described a variety of supports when discussing their switch to ERT. These supports ranged from existing structures that made remote teaching easier to emotional support from colleagues. Here, we describe how we went from transcript to themes (Table 9).

**Table 9.** Themes, categories, and example open codes for supports

| **Example Open Codes** | **Category** | **Theme** |
| --- | --- | --- |
| Access to tools for remote instruction<br>Breakout rooms in Zoom<br>Good internet for instructor | Classroom technology for remote teaching | Tools and support for class content |
| [Mitch] 's remote teaching document<br>Workshops and training from CETL<br>Support from IT with Zoom | Help with technology |  |
| Having a hard-working TA<br>Help from more experienced colleagues<br>Staff support for developing lab material | Help with teaching |  |
| Peer supporter<br>Informal conversations with colleagues<br>Community of colleagues | Community | Mental/emotional support |
| Students understanding/supportive of instructor<br>Flexible mindset<br>Supportive administrative attitude | Socio-emotional factors |  |
| Personal confidence<br>Self-motivation<br>Autonomy was empowering | Teacher beliefs |  |
| Quiet place to work<br>Spouse and pets<br>Able to take a quick nap | Work from home |  |
| IOR created highly structured transition<br>Small changes to accommodate ERT rather than large changes<br>SNS staff made things happen faster | Reducing cognitive load |  |
| Prior knowledge with iPad<br>Prior training about using Canvas well<br>Prior videoconferencing technology knowledge and experience | Prior experience | Preexisting supports |
| Early warning about transition<br>Meeting as a class prior to ERT<br>Timing of switch to ERT | Timing |  |
| Flipped class structure made it easy to switch to remote<br>Having a small class<br>Class structure was easy to transition to remote environment | Course attributes |  |
| Student comfort with online experience<br>Student familiarity with streaming platform<br>Students seemed more comfortable asking questions | Student comfort interacting online |  |

#### Instructor open coding for supports

By starting with open coding, we were able to deductively determine what codes were present in all transcripts. This process allowed us to develop codes that may not have been present in an already formalized coding tool.

An example support is given below:

> *Yeah. So I needed some technical help, just advice about online exams and best practices for doing synchronized learning. But then, the DBER Club and the scientific institutes, those were also just moral support types of help. Just knowing that it was new for everyone, we’re all struggling, and hearing from other folks about particular types of issues to figure out if what I was experiencing was similar or different and, yeah, just getting ideas about how to handle different situations. ---Laney*

During the open coding process CD identified supports that were both technical and moral, while EM identified a support of hearing how others were handling the situation. During the consensus process, discussion between the consensus coders focused on whether to differentiate between technical and moral support, how specific to be with respect to technical support, and whether it was the people or the discussions that were the actual support. This discussion resulted in two codes:

1. Conversations with colleagues about potential ideas and practices
2. Casual conversations with colleagues for moral support

After consensus coding all the supports, a total of 134 unique supports were identified.

#### Instructor categories for supports

After reaching consensus on all the transcripts, the researchers individually collected the codes into categories, then met to resolve any differences. This process led to the creation of 12 categories, shown in Table 9. These categories covered topics, such as access to technology, help with remote teaching pedagogy, access to a supportive community, the comfort of working from home, and the timing of the switch to ERT. As an example of how these support categories came about, consider the two quotes below:

> *I’d say getting the Canvas website very organized has supported the class content delivery. [Claire] ended up looking at my class when I was weaving up the website when all this had to happen—when I was putting this online. So having somebody who had developed detailed Canvas modules before—just gave me a few tips—was really useful right then. ---Constance*
>
> *I felt like, specifically at UC Merced, there was this general community, this sense of community among all of the colleagues and everybody banding together and making it—making the whole learning environment as good as it could be. That’s what it felt like. I felt very fortunate to be at UC Merced working with all these really cool people who are doing the best that they can to make sure that everything’s hunky dory. ---Josephine*

On the surface both quotes seem similar, in that both Constance and Josephine identified people they could talk to as a support. However, Constance’s quote points to the support of having someone available to answer specific, course-related questions about best practices, whereas Josephine’s quote is more about the camaraderie of people in similar situations and knowing that others are persevering under difficult circumstances. As a result of these differences, Constance’s quote was coded as ‘help from more experienced colleagues’, whereas Josephine’s quote was coded as ‘community of colleagues’. These differences also led to the quotes being placed in different categories. As Constance’s quote was focused primarily on pedagogical help, it was placed in the ‘help with teaching’ category, whereas Josephine’s quote, which focused much more on the emotional support of others, was placed in the ‘community’ category.

#### Instructor themes for supports

Once the codes were categorized, the researchers individually organized them into themes, and then met to resolve any differences. This process led to the creation of three broad themes of: 1) tools and support for class content, 2) mental/emotional support, and 3) pre-existing supports, shown in Table 9.

> *So in the classroom, we’ve been using Canvas and Zoom a lot. We’ve been doing breakout sessions, making sure to—mostly with follow-up e-mails a lot, things like that, personally wired headsets, stuff like that, as far as hardware and such like that. I’m not sure if I’m answering the question. ---Chase*

Chase’s quote was representative of the supports that fell into the ‘tools and support for class content’ theme. These were supports those instructors identified as helping them deliver or develop material in a remote environment. Other examples of this theme were workshops from UC Merced’s CETL, online groups that instructors could turn to with pedagogical questions, and the ability to host videos through sites like YouTube.

> *Well, for me, personally, I think the students have been very good, very supportive. Even though I don’t see them anymore, they still have a very good attitude and participate in lectures and worked very hard, take it pretty seriously. So that’s emotionally or psychologically, that’s a very positive factor. ---Yuan*

Yuan’s quote was representative of the supports that fell into the ‘mental/emotional support’ theme, which are supports that help instructors and TAs handle the additional stress and emotional toll of working in a remote environment, under unfamiliar conditions. Other examples of this theme were having a supportive partner at home, enjoying the challenge of trying something new, and having a well-developed transition plan to minimize confusion.

> *One is because of another project I worked on, I was familiar with making videos and editing videos. So the technology was familiar to me. I was already somewhat familiar with Zoom. I’ve already had been doing a fair amount of using CatCourses and Top Hat and web-based tools for teaching alongside my face-to-face. So it wasn’t like I went from totally in class to totally not. I think that helped. ---Danielle*

Danielle’s quote was representative of the supports that fell into the ‘pre-existing supports’ theme. These were supports that were already in place that inadvertently made the transition to remote teaching easier for the instructors. Other examples were instructors who had prior experience with online teaching, student familiarity with streaming services, and courses previously delivered using a flipped modality.

### Instructor barriers

Following supports, we found a larger variety of barriers described by instructors discussing their transition to ERT. These barriers covered issues ranging from instructor access to technology to concerns about academic integrity. Here, we describe how we went from transcript to themes.

#### Instructor open coding for barriers

An example barrier is given below:

> *No, it was a slow decline toward the end of the semester. And you usually get that in classes, anyways, but I was noticing it a whole lot more. Some students just become overwhelmed and slowly stop turning in assignments, but this seemed a little more pronounced than what I’ve usually seen. ---Diane*

During the open coding process CD identified a decrease in student participation as a barrier, while EM identified a decrease in quality of student work as a barrier. During the consensus process, discussion between the consensus coders focused on gauging participation and measuring quality, which resulted in barrier code of ‘decrease in student assignment submissions.’ After consensus coding all the barriers, a total of 203 unique barriers were identified.

#### Instructor categories for barriers

While there were many more barriers than supports coded, the barriers collapsed into a similar number of categories (14 barrier categories, compared to 12 support categories), shown in Table 10. These categories covered topics such as issues with instructors and students using technology, increased responsibility, and workload, decreased student motivation, and concerns about student integrity. As an example of how these barrier categories came about, consider the two quotes, below:

> *With a pre-recorded video, that’s impossible. Then, students won’t even watch it. I was convinced I shouldn’t do asynchronous. Then, through the experience, I discovered that conviction was reinforced, because the students told me. I asked them multiple times, and they told me, “This is much better precisely because we can stop you and just ask you to explain the thing, again.” ---Arturo*
>
> *It’s up to them to use their time the way they want to do it, but what we’re finding is it’s hard to project with the statistics whether they’re actually watching the videos or skimming through the videos or how many times they’re watching the videos. That’s hard data to obtain. The only way to get at it is to maybe get an assignment back from them—a lab or a quiz or whatever. You covered some of the stuff on the videos and kind of points out, “These are some of your flaws that you’re not really looking at the videos.” ---Roberto*

**Table 10.** Themes, categories, and sample codes for barriers

| Example Open Code | Category | Theme |
| --- | --- | --- |
| Harder to engage with students<br>Difficult to encourage peer-peer interactions<br>Hard to communicate math via Zoom | Communication difficulties | Structures getting in the way |
| Delays getting Zoom licenses<br>Instructor access to technology<br>Low instructor internet bandwidth | Instructor technology issues |  |
| IT interference with video creation<br>Institutional pressure to be primary communicator<br>Lack of clear shutdown timeline | Administrative issues |  |
| Student bandwidth limitations<br>Lack of student access to technology<br>Lack of face-to-face technical support for students | Student technical issues |  |
| Having TA grade in Canvas<br>Assessing if students are watching videos<br>Incorporating formative assessment | Assessment difficulties |  |
| Background noise during class time<br>Cooperative and group learning didn't feel feasible<br>Harder to be aware of student issues and motivation | Instructional space |  |
| Covering material more slowly<br>Hard to cover all material by remote instruction<br>Time necessary to develop/modify materials for online labs | Time management | Spending more time and effort |
| Inability to monitor student attention distracted instructor<br>Finding the right balance of pace and flexibility<br>Figuring out how to transition the labs to online format | Instructor teaching inexperience |  |
| Institutional training didn't match instructor's need<br>Lack of staff help to prepare for ERT prior to ERT<br>Difficulty finding necessary resources | Teaching and learning resources |  |
| Managing TAs was challenging<br>Responsibility for adjunct instructors<br>Needing to be in total control of course | Responsibility and workload |  |
| Decreased student accountability<br>Students were less interactive<br>Decrease in office hour attendance | Student presence and participation |  |
| Concerns about academic integrity<br>Hard to prevent cheating in online exams<br>Not being too flexible to avoid being taken advantage of | Student integrity | Affective issues |
| Concerns about student mental health<br>Not knowing the student work environment conditions<br>Student name visibility | Student emotion and comfort |  |
| Frustration with student preparation for lecture<br>Zoom fatigue<br>Difficulties managing work life balance | Instructor emotion |  |

Similar to the support quotes discussed above, both Arturo and Roberto identified barriers that, on the surface, seem similar in that they both present difficulties with video lectures. However, Arturo’s quote really pointed to the difficulty of getting immediate feedback from students about specific topics, whereas Roberto’s quote was more about collecting data about whether students are really watching the videos. As a result of these differences, Arturo’s quote was coded as ‘asynchronous teaching was not possible due to students not being able to provide timely feedback about content understanding’, while Roberto’s quote was coded as ‘assessing if students are watching videos.’ In addition, these differences led to the codes being placed in different categories. Because Arturo’s quote was about the potential difficulty of engaging and interacting with students via an asynchronous modality, his code ended up in the ‘instructional space’ category and Roberto’s quote, which focused much more on difficulties measuring student interaction, ended up in the ‘assessment difficulties’ category.

#### Instructor themes for barriers

Once the codes were categorized, we individually organized them into themes, and then met to resolve any differences. This process led to the creation of three broad themes of: 1) structures getting in the way, 2) spending more time and effort, and 3) affective issues, shown in Table 10. Exemplar quotes for these three themes are shown below.

> *And I think, sometimes, it assumes that what happens in those meetings is being shared with other faculty, and that’s not always the case. And so it would have been nice to hear some of that earlier on. And I think going forward that that should just be made explicitly clear that more information is better, I guess, especially when we’re all separated from each other. So that would have been a nice support to have. ---Laney*

Laney’s quote was representative of the barriers that fell into the ‘structures getting in the way’ theme. These were barriers that instructors identified as preventing them from quickly and efficiently transitioning to ERT. Other examples of this theme were poor bandwidth for instructors, TAs, and students, lack of access to a quiet workspace, and difficulty writing math equations in Zoom.

> *Then I figured out, oh, they can see everybody in the entire class. Then it wouldn’t make it a group discussion. Then I put students in groups where each group is all the students in the discussion session, and the time it took to put that together just sucked. So much of my time got sucked up with this bureaucratic, digital—I had to click buttons. I had to move students. I had to cross-reference things. ---Claire*

Claire’s quote was representative of the barriers that fell into the ‘spending more time and effort’ theme. These were barriers associated with instructors and TAs working less efficiently. Other examples of this theme were having to take time to teach other instructors how to teach remotely, working harder to have students engage with the material, and the inefficiency of communicating via Zoom.

> *I know, and also, my big concerns were for the mental health of my students because they all—my class is really unique, they all live together… and they are all each other’s best friends, and then they just got ripped apart from each other. At the beginning of the week everything was fine, and then that Friday everybody was home and they weren’t seeing each other for the rest of the semester. And I know that that was tough on them. And so I was concerned for everybody’s wellbeing. ---Kiera*

Kiera’s quote was representative of the barriers that fell into the ‘affective issues’ theme. These were barriers associated with instructors or TAs dealing with negative emotions, either theirs or from students. Other examples of this theme were worrying about student academic integrity, witnessing abusive home environments, and feeling guilty about not being an expert in remote teaching.

### Student supports

Students described nine supports when discussing what helped their learning through the transition to ERT, ranging from course attributes to prior experiences that students had before the start of ERT (Table 9). However, ‘course attribute’s (38.6%), ‘technology for remote teaching’ (22.9%), and ‘community’ (11.4%) were the three most frequent categories described by students as supporting their learning during ERT (Figure 4). Here, we describe how we went from student survey responses to categories for Claire’s class. For example, one student wrote that:

> *Homework assignments and discussion sections with my TA were extremely helpful, and the lab sections also helped. ---Sammi*

**Figure 4.**
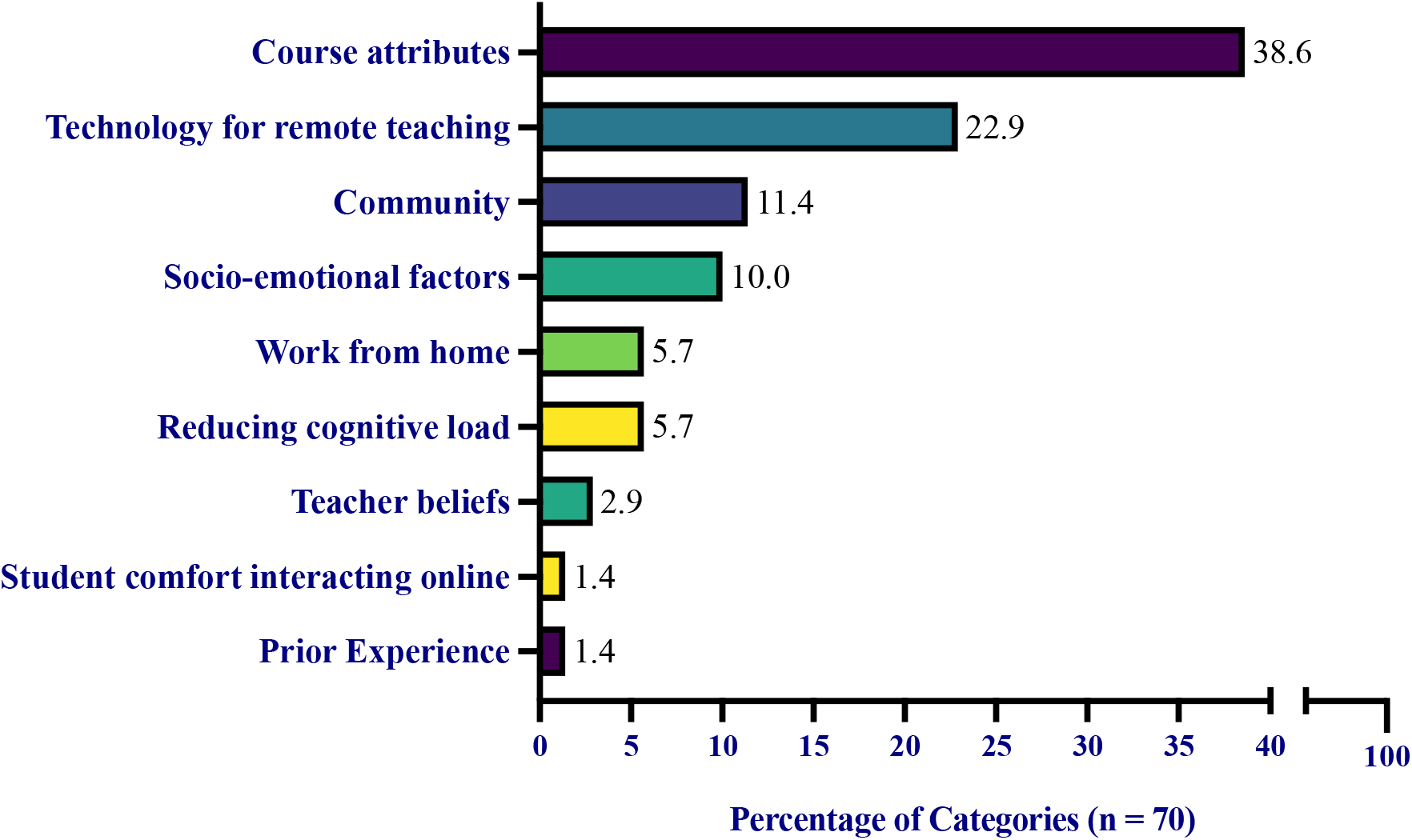
Student perceived supports during their transition to ERT. The percentage of support categories from students. Each value is the total number of each category divided by the total number of all categories, 70. For example, ‘course attributes’ was mentioned 27 times and 27/70 = 38.6%. Note: no student used a single code more than once.

Homework assignments, discussions sections, and lab sections were all attributes of the course that had existed prior to the transition to ERT. Therefore, Sammi’s quote was placed into the category ‘course attributes’ since the student discussed components of the course (e.g., homework assignments) that help them ease into the transition to ERT. This was the most frequent support category described by students. Two other students, Ash and Jax, wrote that:

> *I think the instructional videos provided helped us greatly in learning the content of the course. Zoom calls helped us catch up and know his outlook on the plan for the coming week of the course. ---Ash*
>
> *Zoom discussions, zoom lectures, recorded lectures, online tutoring services outside of school ---Jax*

Ash stated that features of the Zoom platform aided their learning, which lead to their response being put in the category’ classroom technology.’ In addition, Jax listed Zoom lectures, discussions, and other online sources as resources that helped their learning, which lead to this response being placed in the category’ classroom technology.’ Finally, another student wrote that:

> *Having online lectures actually felt like I was more connected to the class rather than in person, so that really helped me out. ---Ekene*

This quote was placed in the category ‘community’ because of the connection to the class that aided the student and helped them sustain their learning.

### Student barrier categories

Students described ten barriers when discussing what challenged their learning during ERT. These barriers ranged from concepts such as instructor technology issues to instructional spaces (Table 10). However, ‘instructional space’ (23.7%), ‘student emotion and comfort’ (21.1%), and ‘student presence and participation’ (21.1%) were the most frequent categories students found challenging their learning during ERT (Figure 5). Here, we described how we went from student survey responses to categories for students from Constance’s class. For example, two students wrote that:

> *That I felt there wasn’t much learned as if I were in person. ---Nasim*
>
> *Not being able to study in groups or ask my friend to help explain my questions. Also, there are fewer physical demonstrations in classes and I tend to get more distracted at home ---Ollie*

**Figure 5.**
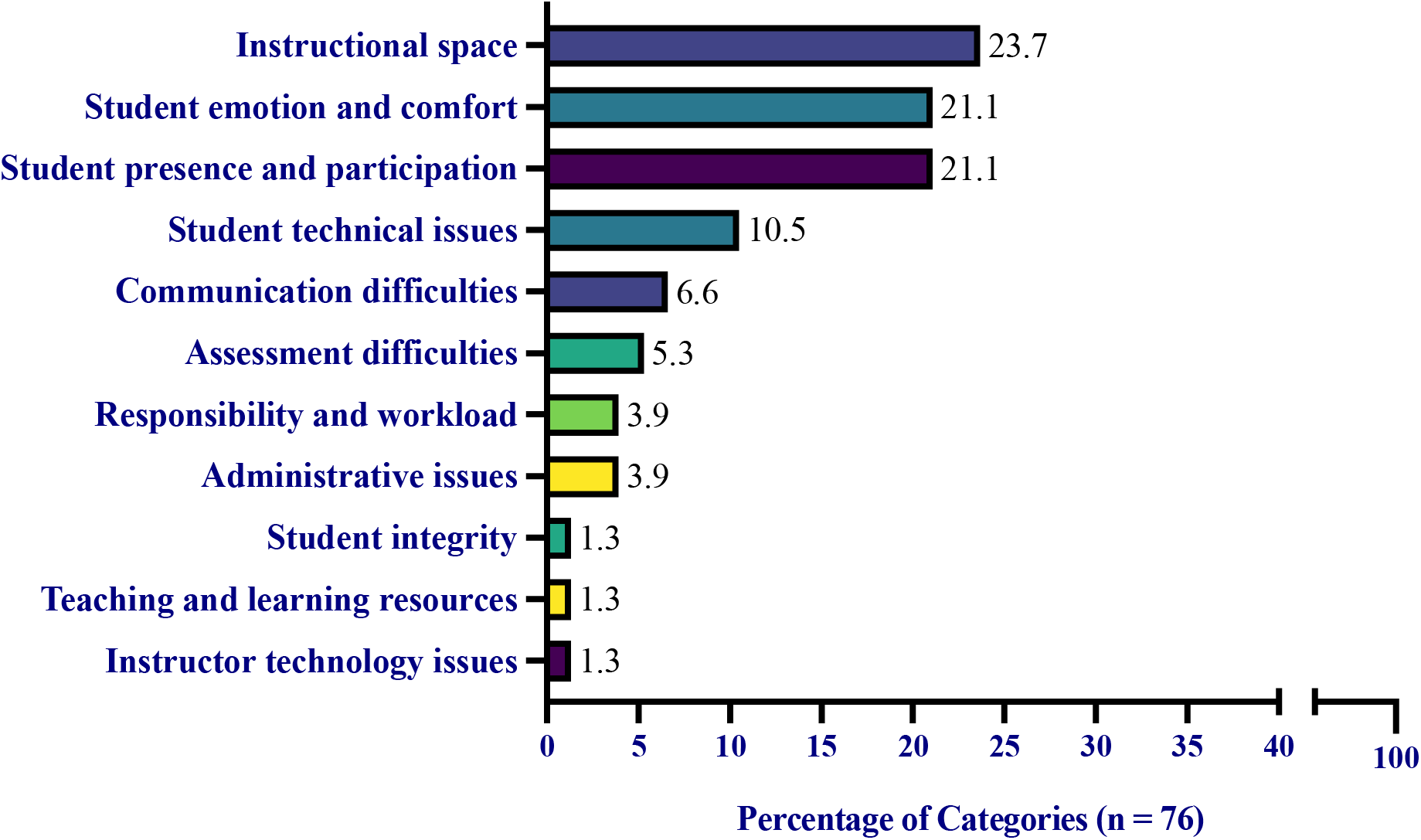
Student perceived barriers during their transition to ERT. The percentage of barriers from students. Each value is the total number of each category divided by the total number of all categories, 76. For example, ‘instructional space’ was mentioned 18 times and 18/76 = 23.7%. Note: no student used a single code more than once.

Nasim described feeling as if they were not learning as much remotely as in-person. Additionally, Ollie described how the remote learning environment was difficult and distracting compared to the in-person learning environment. Therefore, both quotes were put in the category’ instructional space.’ Additionally, two students wrote that:

> *It was hard to focus at home because there were a couple of distractions. Also I kind of lost some motivation through online learning. ---Xia*
>
> *Productivity was a challenge of mine. I live in a household with younger siblings who are not only rowdy but also need help with their online classes as well. ---Rio*

There are many different reasons that might prevent students from attending or participating in course activities or office hours. These factors can range from different emotional states to familial responsibilities and can have a negative impact on students’ learning. Because Xia and Rio described facing some of these difficulties causing them to be unproductive, they were placed in the category’ student presence and participation.’ Lastly, one student wrote that:

> *When we transitioned to [video only] lectures. I felt cut off and sort of isolated from the class. It was starting to feel pretty lonely and discouraging. ---Mitra*

Because Mitra wrote that this isolation was a barrier to their learning, and it resulted in them feeling lonely and discouraged, it was listed as ‘student emotion and comfort.’ We placed it in that category because we felt it could be included as a concern for student emotional well-being.

### Instructor and student comparisons

To further investigate the student experience, we decided to compare the student survey data to their instructors. We wanted to know if the students that we surveyed had a similar experience in their transition to ERT as their instructors. It was fortunate that we had all four instructors of the students surveyed as part of our instructor data. All the student data came from students with at least one course taught by an instructor that also participated in our study, and we wanted to compare the supports and barriers identified by the students with these four instructors. When assessing the instructor and student category data we found that there were more barriers described by students than supports, like the instructors’ results. The most frequent support categories applicable to students were ‘course attributes’, ‘technology for remote teaching’, and ‘community’. In comparison, the instructors of those students’ most frequent support categories were ‘community’, ‘help with teaching,’ and ‘help with technology.’ We found that both instructors and students overlapped with mentioning the ‘community’ category defined as a “personal or professional network to sustain teaching, learning, or yourself.” It is understandable that both students and instructors alike felt community was one of their biggest supports.

> *So by and large, colleagues, and a broader, especially for the courses I teach, a pretty strong community, a professional community outside of the campus. So other specialists in my area that are also engaged in teaching the same kinds of classes ---Haik (Instructor)*
>
> *I like how professor [Instructor Haik] set up a hybrid online course. This allowed me to view the lecture videos on my own time and i still had the opportunity to ask questions and listen to the questions of my peers during zoom lectures. ---Mitra (Student)*

If we look at Haik, the instructor, and one of their students, Mitra, we can see how they both described their perspective colleagues as part of their supportive community. Instructor Haik discussed how his community of support is other specialists in his area of research and Mitra discussed how their peers during synchronous teaching provide support.

The categories’ instructor teaching inexperience’ and ‘time management’ were some of the most common categories from instructors as barriers to their transition to ERT. However, neither of these categories were mentioned by students, showing that even though there may be overlapping supports between students and instructors, the barriers seem to differ more. One common barrier cited by instructors and students was the category’ instructional space’, which we have described as “difficulty implementing or participating in teaching and learning activities.”

> *With breakout rooms, students weren’t really—they needed to be introduced and encouraged, kind of trained, on use. Show your video. Talk to each other. Initiate the conversation. I also found it really difficult that if I was in a breakout room, nobody could contact me. If my TA was like, hey, you need to come back, they couldn’t reach me. ---Claire*
>
> *The difficulty of focusing given the format, and the impersonal-ness of lectures, as well as the difficulty of following lectures when the pace is not as connected with the students’ ability to keep up. ---Sammi*

The instructor Claire and one of her students Sammi both described how ‘instructional space’ inhibited their teaching or learning. Claire described how she found it difficult to get students to work in groups and find the students and groups that needed her help. Sammi felt it was difficult to focus with the format that Claire was using, as well as they felt it was impersonal and did not connected to their pace of learning. It seems that Claire and Sammi both struggled with Zoom, especially the breakout room feature, as the ERT teaching platform.

## DISCUSSION

We contextualized support and barrier categories within the COI framework by illustrating the positive and negative influences on *teaching presence, social presence*, and *cognitive presence* (Figure 2). Our data shows the importance of providing supports to enable instructors and students to maintain strong *teaching, social*, and *cognitive presence*. Without these supports, there are potential impacts on work-life balance and mental health of faculty and students.

### Teaching presence

*Teaching presence* is defined as “the design, facilitation, and direction of cognitive and social processes for the purpose of realizing personally meaningful and educational worthwhile learning outcomes” (Anderson et al., 2001a). One of the themes we considered a support for *teaching presence* was ‘tools and support for class content.’ Having communication tools and the right technology was critical for both students’ and instructor’s success during the transition to ERT. For example, several instructors mentioned how their early access to technology allowed them to quickly transition from in-person to remote. This quick transition helped them maintain their *teaching presence*. In contrast, the instructors that had difficulties with technology, such a delays getting Zoom licenses or poor internet connection fell under the theme ‘structures getting in the way.’ This theme of ‘structures getting in the way’ directly hindered instructors’ ability to provide and maintain a consistent *teaching presence*, thereby hindering student’s ability to experience a strong *teaching presence*.

An example of a support category we considered an influence on *teaching presence* was the category ‘prior experience.’ We defined prior experience as “knowledge, skills, and experiences before the start of ERT.” For example, instructor Diane said: “We did an interview with a Skype scientist, and Zoom worked really well for that, because I was already using Skype, so I just transitioned to that platform,” which indicates her prior knowledge with Skype was a contributing factor to her ability to quickly transition to ERT, and it allowed her to maintain her *teaching presence*, largely unchanged, for that aspect of the course. Additionally, Diane’s prior knowledge was a support in the switch to ERT because it allowed her to maintain meaningful learning outcomes for her students.

In contrast, an example of a barrier category we considered a negative influence on *teaching presence* was ‘instructor’ technology issues.’ This category was defined as “the inability for instructors to have access to hardware and software” and we observed its impact on instructor ability to maintain *teaching presence*. For example, Roberto described how this barrier impacted him:

> *Even with some of the adjustment is—I’ve recorded some of my videos on Zoom and using a whiteboard or whatever. A lot of students are asking me, “Can you write a little better?” “I’m trying to do it a little better, but it’s hard.” Something that would take me to write something on a piece of paper or write something on a board is taking me 10 times longer. It’s 10 times more inefficient mainly because I don’t have the technology at home. ---Roberto*

Unlike Diane, Roberto encountered significant issues with technology in his teaching. He felt that his lack of technology at home created a barrier to providing his students with worthwhile educational experiences. Better access to technology would have allowed Roberto to improve his *teaching presence* for students. Here we see the student Jax having trouble with technology.

> *Unstable internet, unreliable technology, lack of motivation. ---Jax*

They describe how lack of internet and unreliable technology led to their lack of motivation during class.

As we aligned support and barrier categories with *teaching presence, social presence*, and other attributes, we discovered that there were more instances of *teaching presence* supports and barriers than the other two elements of the COI framework. Additionally, knowing that the relationship between barriers and supports is important, we suggest that well developed student *cognitive presence* could lead to an extension of students learning and capabilities (Figure 5).

**Figure 5.**
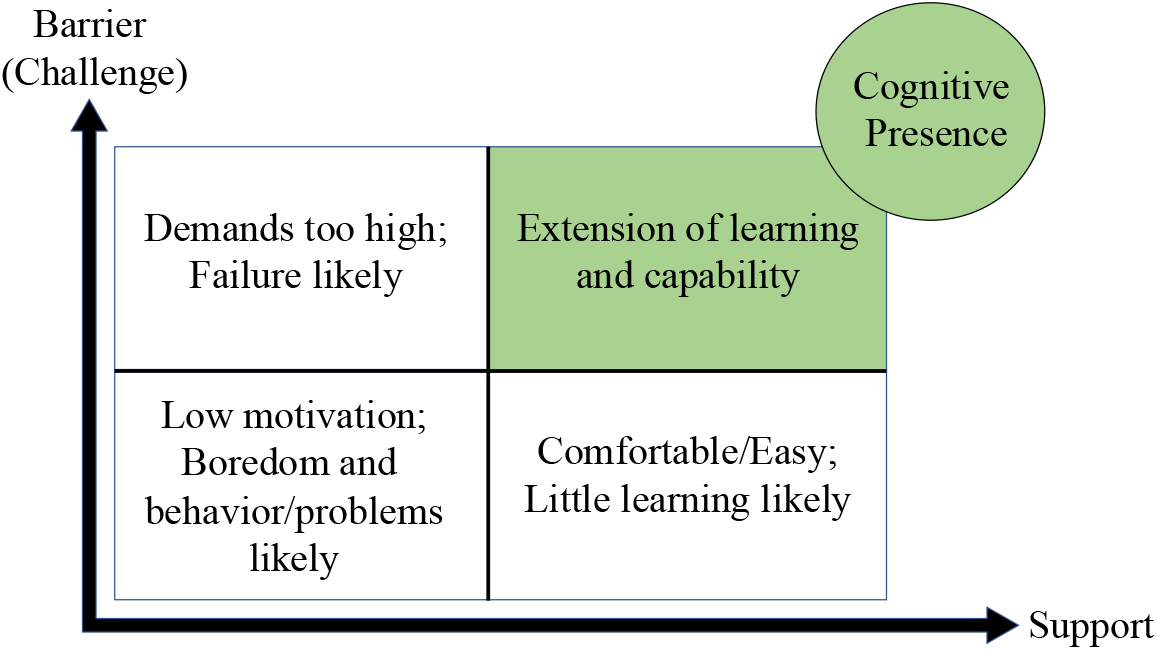
Scaffolding in learning contexts for students. Predicted student outcomes. The top left quadrant, ‘demands too high; failure likely’ is when barriers or challenges are not met with equal support, leading to a likely failure for the student. The lower left quadrant ‘low motivation; boredom and behavior/problems likely’ is generally seen when challenges and supports are both too little. The lower right quadrant ‘comfortable/easy; little learning likely’ is when challenges are too low but supports are high. The last quadrant, the upper right is when challenges or barriers and supports are equal leading to an ‘extension of learning and capability’ Adapted from Hammond & Gibbons (2005).

### Social presence

*Social presence* is “the ability of learners to project their personal characteristics into the community of inquiry, thereby presenting themselves as ‘real people’”. The ‘community’ category within the theme of ‘mental and emotional support’ was something both students and instructors mentioned as important in their transition to ERT. Instructors discussed things like having a support group to allow them to learn from others and having friends to vent to about their frustrations. Having a sense of community provided a basis for *social presence* in that in our study we considered instructors as learners as well as students. Instructors in ERT were learning how to teach remotely in a pandemic; therefore, instructors’ *social presence* was also something important to consider in our analysis.

Another category that aligned with *social presence* was ‘student comfort interacting online’, which was defined as “using communication tools and modalities familiar to students.”

> *I found that the students were very engaged. It didn’t really puzzle them, this remote instruction to online. They were already prepared for it in other classes and the students are all doing very well in this class. ---Aarush*

Instructor Aarush described that his students’ familiarity with online classes allowed them to be more engaged, and thereby increased their *social presence*. However, other instructors saw a decrease in student *social presence* as the semester progressed:

> *And then I guess, yeah, towards the end or towards the middle and the end of the semester, we started seeing less participation, except for those few students that were still engaged. Yeah. So I don’t know. It was crazy because, in the beginning, we’re like, all right, yeah, we got this kind of thing. But then at the end, people were like, ah. ---Katherine*

Instructor Katherine’s students’ enthusiasm and engagement decreased towards the end of the semester. This barrier to ERT was categorized as ‘student presence and participation,’ which was defined as the “inability of students to attend or participate in class, office hours, or course activities.” We argued that lack of student presence and participation decreases student’s *social presence*.

### Cognitive presence

*Cognitive presence* is defined as “the extent to which learners can construct and confirm meaning through sustained reflection and discourse.” The support theme ‘mental and emotional support’ was important because poor mental health and distractions impacted motivation to explore course content, the first component of *cognitive presence*. For example, the support category ‘work from home’ allowed for personal support from spouses and pets, didn’t clearly indicate that there was an impact on *teaching* or *social presence*, but it clearly had an impact on instructors’ ability to teach during ERT.

For example, instructor Martin described how working from home allowed him to feel supported, “And, of course, informal support in mental issues, well, my spouse and my pets were really imperative to keep me sane.” Despite this category not pertaining to *teaching* or *social presence*, it is still particularly important in describing experiences during the transition to ERT. Mental health, critical to both instructor and student success, has been shown to play a significant role in the lives of many during the COVID-19 pandemic (Copeland et al., 2021; Kecojevic et al., 2020; Roman, 2020).

Many instructors found it difficult to manage a proper work-life balance as shown in the barrier category ‘instructor emotion,’ which was defined as “frustration, fatigue, and guilt due to remote delivery.” Instructor Claire described, “There were multiple days where, if I did shower, it was at 3 in the afternoon and I was sobbing in the shower because I just couldn’t see it getting any better, and still try to maintain quality.” Claire’s struggle to maintain a separation between work and home life, coupled with her feelings of hopelessness, did not align with *teaching* or *social presence*, but might have impacted how she engaged with ERT.

## CONCLUSIONS

In this article, we have examined the perceived supports and barriers that affected instructors and students during the rapid transition to ERT resulting from the COVID-19 pandemic. A total of 31 STEM instructors were interviewed about supports and barriers that they had experienced during the ERT transition, and interview transcripts were analyzed via two-cycle quantitative analysis process drawing from grounded theory methods. This process led to the identification of 134 unique supports that were collected into 12 categories, best represented by the themes 1) Tools and support for class content; 2) Mental/emotional support; and 3) Pre-existing support. Similarly, instructors described 203 unique barriers that we collected into 14 categories, best represented by the themes 1) Tools and support for class; 2) Spending more time and effort; and 3) Affective issues.

At the same time, 69 undergraduate students in STEM classes were surveyed about supports and barriers that they had experienced during the ERT transition. The student survey responses were analyzed by inductive coding, using the codes identified during the instructor analysis. The most important support categories identified by students were ‘course attributes’, followed by ‘technology for remote teaching’ and ‘community’. The biggest barriers identified by students fell into the ‘instructional space’ category, followed by the ‘student emotion and comfort’ category and the ‘student presence and participation’ category.

Considering these supports and barriers in the COI framework, we found that some faculty found ways to maintain a strong *teaching presence* and a healthy work-life balance, but most of these supports were local and existed prior to the transition. Despite institutional efforts most of the broad, large-scale supports that faculty identified were logistical in nature and primarily helped the faculty to maintain a strong *teaching presence*. While faculty may not be familiar with the COI framework, they all recognized the importance of maintaining a strong *teaching presence* and facilitating a strong *social presence* for students. However, most of the faculty identified several barriers that prevented them from maintain and facilitating *teaching* and *social presences*. In addition, most instructors had trouble maintaining a healthy work-life balance and found the transition to be extremely stressful. All of this suggests that either 1) the supports provided by the institution were not meeting the instructor and student needs or 2) the instructors and students were unaware of institutional supports. None of the instructors or students identified institutional supports that helped them maintain a healthy work-life balance nor supported them in maintaining a strong *social presence* during ERT. Following this, recommendations for future emergencies for universities could include either more supports for instructors and students, such as greater access to free technology, both software and hardware as well as increased access to reliable internet bandwidth, or, finding better ways to disseminate where and how to get these supports. Additionally, maintaining a healthy work-life balance is vital for everyone. Increased workload was cited by many students and instructors as a significant struggle they faced. It is crucial that universities find ways to help alleviate this added strain, whether it be through hiring additional staff to create things like laboratory instructional videos or simply lowering expectations so that instructors and students that have family obligations directly stemming from the pandemic can attend to them. Additional time for transitions for preparation of new online course material or training for students to become more familiar with remote platforms would also be helpful.

### Limitations and future directions

We acknowledge that there are several factors that limit our study; however, these limitations provide opportunities for future studies. First, we conducted a convenience sample at only one MSI, UC Merced, so there is limited generalizability. Also, we did not employ a systemic approach to ensure even distribution of faculty and students across STEM disciplines at our institutions, so there were more chemistry and biology participants based on the departments of the co-authors. In particular, the students surveyed were from only four of the 31 instructors interviewed. As the transition to ERT during the COVID-19 pandemic was a one-time event, we will not be able to collect more data from other MSIs or within UC Merced to see if our patterns persist in different study contexts or with more participants. It might be possible to work with other MSI’s that conducted similar work and compare our survey data for patterns. Second, this study focuses on self-report survey and interview data that is often perceived as less objective compared to well-developed and validated classroom observation protocols (AAAS 2013). A risk of surveys and interviews is that the participants inaccurately self-report their teaching and learning experiences, limiting the conclusions that can be made from these data. Our future work aims to triangulate the self-report survey and interview data with classroom observations and student learning gains data (i.e., concept inventories) to better understand classrooms interactions and student learning. Third, this study only described the perceived supports and barriers through the transition to ERT, not the continuation of it. Therefore, we collected instructor interview data at the end of the Fall 2020 semester to better understand what served as supports and barriers for instructors during the continuation of ERT. Taken together, we found there are more barriers than supports and that increased resources and communication between students, TAs, staff, faculty, and administration could help mitigate future emergency remote hardships and experiences.

## Acknowledgements

First, we would like to thank the instructors and students who participated in our research, without them this work would not have been possible. We would also like to thank our collaborators at the University of Minnesota in the Schuchardt and Warfa research groups for their valuable feedback in validating our categories, themes, and definitions. Additionally, we would like to acknowledge the UCM IRB team as well as all the members of the Kranzfelder lab and the discipline-based education research journal club at UCM.

## Author contributions

PK and EM acquired funding and supervised the research team. CD, HB, PK, and EM contributed to the study conception and design. Material preparation and data collection were performed by CD, JA, EM, and PK. Data analysis was performed by CD, HB, MCK, EM, and PK. The first draft of the manuscript was written by primarily by CD, with assistance from PK, EM, and HB. All authors commented on previous versions of the manuscript. All authors read and approved the final manuscript.

