## Supplemental Materials for "I will teach you here or there, I will try to teach you anywhere: perceived supports and barriers for emergency remote teaching during COVID-19 pandemic"

**Supplemental Table 1**. Support themes, categories, category descriptions and example codes presented to expert feedback panel

| Theme | Category | Description | Example codes |
|---|---|---|---|
| **Tools and support for class content** | **Prior Experience** | Knowledge, skills, and experiences instructor had before the start of ERT | Prior knowledge with iPad |
| | | | Prior training about using Canvas well |
| | | | Prior videoconferencing technology knowledge and experience |
| | **Timing** | Timing of campus switch to ERT and/or knowledge of when ERT switch was going to happen and the ability to respond | Early warning about transition |
| | | | Meeting as a class prior to ERI |
| | | | Move to ERI happened after students gained necessary hand-on lab experience |
| | **Course attributes** | Components of the course and instructional approaches eased transition | Flipped class structure made it easy to switch to remote |
| | | | Having a small class |
| | | | Class was already broken up in teams, making it easy to utilize breakout rooms |
| | **Student comfort interacting online** | Using communication tools and modalities familiar to students | Student comfort with online experience |
| | | | Student familiarity with streaming platform |
| | | | Students seemed more comfortable asking questions |
| **Preexisting supports** | **Classroom technology** | The ability for students and instructors to access hardware, software, and instructional resources | Access to tools for remote instruction |
| | | | Breakout rooms in zoom |
| | | | Good internet for instructor |
| | **Help with technology** | Advice and assistance using hardware and software | Mitch Calvin's remote teaching document |
| | | | Workshops and training from CETL |
| | | | Support from IT with Zoom |
| | **Help with teaching** | Advice and assistance using online pedagogies | Having a hard-working TA |
| | | | Help from more experienced colleagues |
| | | | Staff support for developing lab material |
| **Mental and emotional support** | **Community** | Personal or professional network to sustain teaching or yourself | Peer supporter |
| | | | Informal conversations with colleagues |
| | | | Community of colleagues |
| | **Affect** | Understanding and affirmation between instructors, students, and administration | Students understanding/supportive of instructor |
| | | | Flexible mindset |
| | | | Supportive administrative attitude |
| | **Self-efficacy** | Positive outlook and confidence in self to succeed | Personal confidence |
| | | | Self-motivation |
| | | | Willing and excited to make changes and try new things-Finds new opportunities invigorating |
| | **Comfort of working from home** | Convenience of working in a comfortable environment | Quiet place to work |
| | | | Spouse and pets |
| | | | Able to take a quick nap |
| | **Reducing cognitive load** | Logistical changes to make transition easy for oneself | IOR created highly structured transition |
| | | | Small changes to accommodate ERI rather than large changes for online teaching |
| | | | SNS staff made things happen faster |

**Supplemental Table 2**. Barrier themes, categories, category descriptions and example codes presented to expert feedback panel

| Theme | Category | Description | Example codes |
|---|---|---|---|
| Structures getting in the way | **Communication difficulties** | Hard to transmit information and resources, inability to read body language, and lack of tone transmission through electronic interactions | Ambiguity with student communication using chat - hard to tell who's talking to whom |
| | | | Difficult to encourage peer-peer interactions |
| | | | Hard to communicate math via Zoom |
| | **Instructor technology issues** | Inability for instructors to access hardware and software | Delays getting Zoom licenses |
| | | | Instructor access to technology |
| | | | Low instructor internet bandwidth |
| | **Student technical issues** | Inability for students to access hardware and software | Student bandwidth limitations |
| | | | Lack of student access to technology |
| | | | Lack of face-to-face technical support for students |
| | **Administrative issues** | Lack of or inconsistent guidance and communication from leadership | IT interference with video creation |
| | | | Institutional pressure to be primary communicator |
| | | | Lack of clear shutdown timeline |
| | **Assessment difficulties** | Lack of student learning and inability to measure it in an online environment | Having TA grade in Canvas |
| | | | Assessing if students are watching videos |
| | | | Incorporating formative assessment |
| | **Classroom environment** | Difficulty implementing teaching and learning activities | Background noise during class time |
| | | | Cooperative and group learning didn't feel feasible |
| | | | Harder to be aware of student issues and motivation |
| Spending more time and effort | **Time management** | Increased time on items other than content | Covering material more slowly |
| | | | Hard to cover all material by remote instruction |
| | | | Time necessary to develop/modify materials for online labs |
| | **Instructor teaching inexperience** | Lack of knowledge and best practices on how to deliver instruction remotely | Inability to monitor student attention distracted instructor |
| | | | Finding the right balance of pace and flexibility |
| | | | Figuring out how to transition the labs to online format |
| | **Teaching and learning resources** | Hard time finding or lack of advice and assistance for using online pedagogies | Institutional training didn't match instructor's need |
| | | | Lack of staff help to prepare for ERI prior to ERI |
| | | | Difficulty finding necessary resources |
| | **Increased responsibility and workload** | Having more things to do because of the transition | Managing TAs was challenging |
| | | | Having to teach colleagues how to save time teaching remotely with tips and tricks |
| | | | Needing to be in total control of course |
| | **Student availability** | Inability of students to attend or participate in class, office hours, or course activities | Decreased student accountability |
| | | | Students were less interactive |
| | | | Decrease in office hour attendance |
| Affective issues | **Student emotion and comfort** | Concern about student safety, health, and happiness | Concerns about student mental health |
| | | | Not knowing the student work environment conditions |
| | | | Student name visibility |
| | **Student integrity** | Assumption that instructors are being taken advantage of or that students are going to benefit by not being fair and honest | Concerns about academic integrity |
| | | | Discomfort using lock-down browser to prevent cheating on exam |
| | | | Not being too flexible to avoid being taken advantage of |
| | **Instructor emotion** | Frustration, fatigue, and guilt due to remote delivery | Clash between content and the real world impacts instructor motivation |
| | | | Instructor feeling guilty about not being able to provide learning experiences they want to |
| | | | Difficulties managing work life balance |

**Supplemental Table 3**. Expert feedback panel answers for supports from Group 1

**Supports- Group 1**

| Quote | Group Category Answers | Group Notes | Correct Answer |
|---|---|---|---|
| That's probably a white male privilege thing, too. I was not too bothered. I have been not too bothered about trying to work standing up in my kitchen except for when there's just a little too much Simpsons and screaming in the background. Then I start to lose it. **We don't really have a home office, but we were lucky that we converted our garage to a sound-isolated music room.** We were a little lucky about the timing of that, because we had just finished that. But it would seem like at least California, if not the country, should maybe invest in broadband access. | Comfort of Working Home | Because it's talking about a comfortable space, don't see other category that works better. "Comfort of Working from Home" was a bit confusing because of the working from home piece. | Comfort of Working Home |
| **One is because of another project I worked on, I was familiar with making videos and editing videos. So the technology was familiar to me. I was already somewhat familiar with Zoom. I've already had been doing a fair amount of using CatCourses and Top Hat and web-based tools for teaching alongside my face-to-face. So it wasn't like I went from totally in class to totally not. I think that helped.** | Prior Experience | It's talking about prior experience with the technologies. Should there be a distinction between technologies and class content and pedagogy? | Prior Experience |
| Well, I guess getting the Zoom installed pretty quickly because I didn't have that installed or an account for that. But also, I guess **what helped was, I don't know, I guess having the tools necessary. Because I know some TAs didn't have a touchscreen laptop or whatever, so it made it more difficult to teach. I mean, I had my tablet that I could use, so I feel like that made it really helpful. Because then, I could just—I also recorded the lecture so I can share it with them.** | Classroom Technology | They had the technology available to help with remote learning. Maybe remote learning technology or technology for remote teaching because no longer in classroom. | Classroom Technology |

* **Bold text** is the text used to define the category

**Supplemental Table 4**. Expert feedback panel answers for supports from Group 2

**Supports- Group 2**

| Quote | Group Category Answers | Group Notes | Correct Answer |
|---|---|---|---|
| That's probably a white male privilege thing, too. I was not too bothered. I have been not too bothered about trying to work standing up in my kitchen except for when there's just a little too much Simpsons and screaming in the background. Then I start to lose it. **We don't really have a home office, but we were lucky that we converted our garage to a sound-isolated music room.** We were a little lucky about the timing of that, because we had just finished that. But it would seem like at least California, if not the country, should maybe invest in broadband access. | Comfort of Working Home | Not technology related, not about classroom stuff, working from home fits best | Comfort of Working Home |
| **One is because of another project I worked on, I was familiar with making videos and editing videos. So the technology was familiar to me. I was already somewhat familiar with Zoom. I've already had been doing a fair amount of using CatCourses and Top Hat and web-based tools for teaching alongside my face-to-face. So it wasn't like I went from totally in class to totally not. I think that helped.** | Prior Experience | Is this also access to technology? But they aren't centering the access rather they are centering the knowledge so we went with that one. | Prior Experience |
| Well, I guess getting the Zoom installed pretty quickly because I didn't have that installed or an account for that. But also, I guess **what helped was, I don't know, I guess having the tools necessary. Because I know some TAs didn't have a touchscreen laptop or whatever, so it made it more difficult to teach. I mean, I had my tablet that I could use, so I feel like that made it really helpful. Because then, I could just—I also recorded the lecture so I can share it with them.** | Classroom Technology | We assumed that the codes were based around the interviewee receiving help or etc not if they were giving help to other instructors which left us not using help with technology | Classroom Technology |

* **Bold text** is the text used to define the category

**Supplemental Table 5**. Expert feedback panel answers for barriers from Group 1

**Barriers- Group 1**

| **Quote** | **Group Category Answers** | **Group Notes** | **Correct Answer** |
|---|---|---|---|
| Yes. And this is another fact. **A lot of them don't want to talk, because they have a background ambiance that they don't want to show to you. They might be ashamed you might hear shouts, noises, or even if they are very low-income and the house is very modest, they don't want to show it.** I don't know. | Trouble deciding whether it's both Student Availability and Student Emotion and Comfort | It wasn't clear whether student emotion and comfort had to be expressed explicitly by the instructor or whether just noticing the conditions might be sufficient. | Student emotion and comfort |
| No, it was a slow decline toward the end of the semester. And you usually get that in classes, anyways, but I was noticing it a whole lot more. Some students just become overwhelm and **slowly stop turning in assignments, but this seemed a little more pronounced than what I've usually seen.** | Trouble deciding whether it's both Student Availability and Student Emotion and Comfort | It seems like both. The availability to do class tasks is impacted, but it's impacted because of the extra effort required throughout the semester. The instructor is noticing an emotional state of the students but they're not expressing a concern for their well-being. | Student availability |
| **With a pre-recorded video, that's impossible. Then, students won't even watch it. I was convinced I shouldn't' do asynchronous. Then, through the experience, I discovered that conviction was reinforced, because the students told me. I asked them multiple times, and they told me, "This is much better precisely because we can stop you and just ask you to explain the thing, again."** | Communication difficulties | | Classroom environment |

* **Bold text** is the text used to define the category

**Supplemental Table 6**. Expert feedback panel answers for barriers from Group 2

| **Barriers- Group 2** | | | |
|---|---|---|---|
| **Quote** | **Group Category Answers** | **Group Notes** | **Correct Answer** |
| Yes. And this is another fact. **A lot of them don't want to talk, because they have a background ambiance that they don't want to show to you. They might be ashamed you might hear shouts, noises, or even if they are very low-income and the house is very modest, they don't want to show it.** I don't know. | Student availability | Maybe technical stuff? Also might be about not being able to assess (no see students through video) | Student emotion and comfort |
| No, it was a slow decline toward the end of the semester. And you usually get that in classes, anyways, but I was noticing it a whole lot more. Some students just become overwhelm and **slowly stop turning in assignments, but this seemed a little more pronounced than what I've usually seen.** | Student emotion and comfort | Maybe also a focus on not being able to assess properly with lack of assignments? | Student availability |
| **With a pre-recorded video, that's impossible. Then, students won't even watch it. I was convinced I shouldn't' do asynchronous. Then, through the experience, I discovered that conviction was reinforced, because the students told me. I asked them multiple times, and they told me, "This is much better precisely because we can stop you and just ask you to explain the thing, again."** | Classroom environment | Note: classroom environment category doesn't match with description; chose this one maybe because it's the broadest | Classroom environment |

* **Bold text** is the text used to define the category

**Supplemental Table 7**. Support themes, categories and their descriptions

**Supports**

| **Theme** | **Category (Version 1)** | **Description (Version 1)** | **Category (Final)** | **Description (Final)** |
|---|---|---|---|---|
| **Tools and supports for class content** | Prior Experience | Knowledge, skills, and experiences instructor had before the start of ERT | Prior Experience | Knowledge, skills, and experiences before the start of ERT |
| | Timing | Timing of campus switch to ERT and/or knowledge of when ERT switch was going to happen and power to make changes in response | Timing | Timing of campus switch to ERT and/or knowledge of when ERT switch was going to happen and the ability to respond |
| | Preexisting course structure | Attributes of the course and instructional approaches eased transition | Course attributes | Components of the course and instructional approaches eased transition |
| | Student comfort | Using communication tools and modalities familiar to students | Student comfort interacting online | Using communication tools and modalities familiar to students |
| **Preexisting supports** | Classroom Technology | The ability for students and instructors to access hardware and software | Technology for remote teaching | The ability to access hardware, software, and instructional resources |
| | Help with technology | Advice and assistance using hardware and software | Help with technology | Advice and assistance using hardware and software |
| | Help with teaching | Advice and assistance using online pedagogies | Help with teaching | Advice and assistance using online pedagogies |
| **Mental and emotional support** | Community | Personal or professional network to sustain teaching or yourself | Community | Personal or professional network to sustain teaching, learning or yourself |
| | Affect | Understanding and affirmation between instructors, students, and administration | Socio-emotional factors | Common understanding and affirmation between instructors, students, and administration |
| | Rising to the challenge | Self-efficacy and positive outlook | Teacher beliefs | Positive outlook and confidence in self to succeed |
| | Comfort of working from home | Convenience of working in a comfortable environment | Work from home | Convenience of working in a comfortable environment |
| | Minimizing strain | Logistical changes to make transition easy | Reducing cognitive load | Logistical changes to make transition easy for oneself |

**Supplemental Table 8**. Barrier themes, categories and their descriptions

| Barriers | | | | |
|---|---|---|---|---|
| **Theme** | **Category (Version 1)** | **Description (Version 1)** | **Category (Final)** | **Description (Final)** |
| **Structures getting in the way** | Communication difficulties | Hard to transmit information and resources, inability to read body language. Lack of tone transmission through electronic interactions | Communication difficulties | Hard to transmit information and resources, inability to read body language, and lack of tone transmission through electronic interactions |
| | Instructor technology issues | Inability for instructors to access hardware and software | Instructor technology issues | Inability for instructors to access hardware and software |
| | Student technical issues | Inability for students to access hardware and software | Student technical issues | Inability for students to access hardware and software |
| | Administrative issues | Lack of or inconsistent guidance and communication from leadership | Administrative issues | Lack of or inconsistent guidance and communication from leadership |
| | Assessment difficulties | Inability to measure student learning in an online environment | Assessment difficulties | Lack of student learning and inability to measure it in an online environment |
| | Classroom environment | Concern about student interaction and learning during class time | Instructional space | Difficulty implementing or participating in teaching and learning activities |
| **Spending more time and effort** | Time management | Increased time administrating and on non-content led to decreased content | Time management | Increased time on items other than content |
| | Instructor teaching inexperience | Lack of knowledge and best practices on how to deliver instruction remotely | Instructor teaching inexperience | Lack of knowledge and best practices on how to deliver instruction remotely |
| | Teaching resources | Hard time finding or lack of advice and assistance for using online pedagogies | Teaching and learning resources | Hard time finding or lack of advice and assistance for using online pedagogies |
| | Increased responsibility and workload | Having more things to do because of the transition | Responsibility and workload | Increased cognitive effort from having to develop new habits and techniques |
| | Student availability | The general desire or willingness of students to attend class, office hours or other course activities | Student presence and participation | Inability of students to attend or participate in class, office hours, or course activities |
| **Affective Issues** | Student emotion and comfort | Concern about student safety, health, and well-being | Student emotion and comfort | Concern about safety, health, and emotional well being |
| | Student integrity | Assumption that instructors are being taken advantage of or that students are going to benefit by not being fair and honest | Student integrity | Concern that instructors are being taken advantage of or students are going to benefit by not being fair and honest |
| | Instructor emotion | Frustration, fatigue, and guilt due to remote delivery | Instructor emotion | Frustration, fatigue, and guilt due to remote delivery |
